# The Antibiotic Neomycin Enhances Coxsackievirus Plaque Formation

**DOI:** 10.1101/471284

**Authors:** Mikal A. Woods Acevedo, Julie K. Pfeiffer

## Abstract

Coxsackievirus typically infects humans via the gastrointestinal tract, which has a large number of microorganisms collectively referred to as the microbiota. To study how the intestinal microbiota influence enteric virus infection, several groups have used an antibiotic regimen in mice to deplete bacteria. These studies have shown that bacteria promote infection with several enteric viruses. However, very little is known about whether antibiotics influence viruses in a microbiota-independent manner. Here, we sought to determine the effects of antibiotics on coxsackievirus B3 (CVB3) using an *in vitro* cell culture model in the absence of bacteria. We determined that an aminoglycoside antibiotic, neomycin, enhanced plaque size of CVB3-Nancy strain. Neomycin treatment did not alter viral attachment, translation, or replication. However, we found that the positive charge of neomycin and other positively charged compounds enhanced viral diffusion by overcoming the negative inhibitory effect of sulfated polysaccharides present in agar overlays. Overall, these data lend further evidence that antibiotics can play non-canonical roles in viral infections and that this should be considered when studying enteric virus-microbiota interactions.

**Importance:** Coxsackieviruses primarily infect the gastrointestinal tract of humans, but they can disseminate systemically and cause severe disease. Using antibiotic treatment regimens to deplete intestinal microbes in mice, several groups have shown the bacteria promote infection with a variety of enteric viruses. However, it is possible that antibiotics have microbiota-independent effects on viruses. Here, we show that an aminoglycoside antibiotic, neomycin, can influence quantification of coxsackievirus in cultured cells in absence of bacteria.

## Introduction

Coxsackievirus B3 (CVB3) is a cardiotropic nonenveloped RNA virus belonging to the *Enterovirus* genus of the *Picornaviridae* family. CVB3 is an important human pathogen, which can cause a wide range of diseases, including myocarditis, cardiac arrhythmias, aseptic meningitis, type 1 diabetes, gastrointestinal distress, and death (1–5). CVB3 has been implicated in over 40,000 infections a year in the United States alone and there are no current treatments or vaccines for CVB3 infections (6).

Within the gastrointestinal tract resides a microbial ecosystem of approximately 10^14^ organisms, which play a crucial role in host homeostasis (7). The intestinal microbiota can also influence infection with orally acquired enteric viruses (8–10). Alterations in microbiota, for example through antibiotic treatment, can influence enteric pathogen susceptibility (8–10). However, not much is known about direct effects of antibiotics on enteric viruses.

Antibiotics can have a variety of microbiota-independent effects on mammalian cells. Antibiotics can illicit profound changes in host gene expression in both conventional and germ-free mice (11), alter mammalian metabolic pathways and impair the phagocytic activity of immune cells (12), induce mitochondrial dysfunction (13, 14), and inhibit histone demethylases (15). Additionally, Gopinath et al. recently demonstrated that aminoglycoside antibiotics can confer microbiota-independent antiviral resistance against both DNA and RNA viruses by upregulating expression of interferon-stimulated genes (16).

Here, we examined the effect of antibiotic treatment on CVB3 infection of cultured cells in the absence of bacteria. From a group of antibiotics that is commonly given to mice in microbiota depletion studies, we found that neomycin increases plaque size of CVB3. Notably, treatment with neomycin did not have an apparent effect on viral replication in single cycle growth curves. We determined that plaque size enhancement by neomycin was most likely due its positive charge overcoming the inhibitory negative charge of agar overlays, thus aiding viral diffusion.

## Results

### Neomycin increases plaque size of CVB3-Nancy, but not poliovirus

To examine the effect of antibiotics on plaque formation of CVB3-Nancy, we infected a monolayer of HeLa cells that were pretreated with or without 1 mg/ml of an antibiotic cocktail consisting of vancomycin, ampicillin, neomycin, and streptomycin. Following adsorption for 30 min, inoculum was removed, and an agar overlay with or without antibiotics was added. To visualize plaques, plates were stained with crystal violet 3 days post infection. When cells were exposed to the antibiotic cocktail, we observed a significant increase in CVB3-Nancy plaque size (Fig. 1A). Treatment with vancomycin, ampicillin, or streptomycin alone did not confer the large plaque phenotype (Fig. 1A), but treatment with neomycin was sufficient for the large plaque phenotype (Fig. 1B). We quantified plaque size and found that when cells were exposed to neomycin, CVB3-Nancy plaques averaged 6.9 mm^2^, while CVB3-Nancy plaques in untreated cells averaged 0.11 mm^2^ (Fig. 1C). We next determined whether neomycin also affects the plaque size of a closely related enteric virus, poliovirus. When cells were pretreated with or without neomycin and infected with poliovirus, plaques were relatively large and no increase in plaque size was observed with neomycin treatment (Fig. 1D and 1E). Overall, these data indicate that treatment with neomycin is capable of increasing plaque size CVB3-Nancy, but not poliovirus.

**Figure 1.**
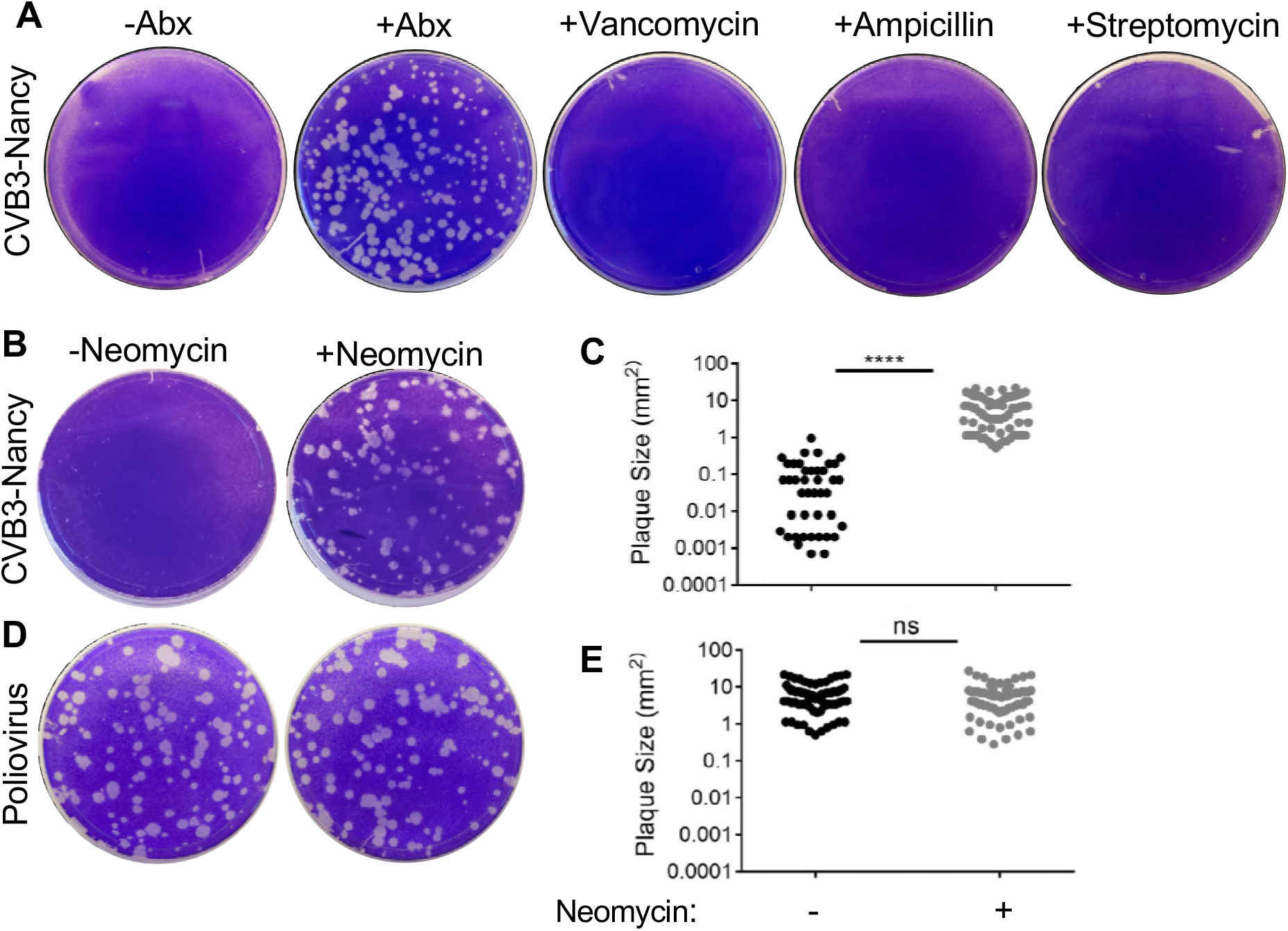
Effect of neomycin on plaque formation of CVB3-Nancy and poliovirus. A) Effects of antibiotics on CVB3-Nancy plaque formation. HeLa cells were pretreated with or without 1 mg/ml of the indicated antibiotics prior to plating 100 PFU of CVB3-Nancy on cells with agar overlays with or without 1 mg/ml of each antibiotic. Plates were stained with crystal violet 48 hpi. Abx= ampicillin, neomycin, streptomycin, and vancomycin mixture. Effects of neomycin on CVB3-Nancy (B) or poliovirus (D). C) Plaque size quantification of B. E) Plaque size quantification of D. Each symbol represents a plaque. ****, P<0.0001 (unpaired two-tailed Student *t* tests). ns= not significant.

### Effect of neomycin on plaque size of different CVB3 strains

Since neomycin was capable of increasing plaque size of CVB3-Nancy, but not poliovirus, we hypothesized that neomycin may also increase plaque size of closely related CVB3 strains. To investigate this hypothesis, we first used CVB3-Nancy-N63Y, which is a CVB3-Nancy derivative that contains a single point mutation in the VP3 capsid protein, N63Y, which induces formation of large plaques in agar overlays due to reduced binding to sulfated glycans (17). We found that neomycin was also capable of increasing plaque size of CVB3-Nancy-N63Y in agar overlays (Fig. 2), although the effect was less pronounced when compared to CVB3-Nancy due to the larger plaques of CVB3-Nancy-N63Y in untreated cells. We next examined if neomycin treatment could increase the plaque size of CVB3-H3, a strain of CVB3 that is more virulent in mice (18). Neomycin treatment did not alter CVB3-H3 plaque size (Fig. 2). These data indicate that in agar overlays neomycin is capable of increasing the size of CVB3-Nancy and a closely related mutant CVB3-Nancy-N63Y, but not of CVB3-H3.

**Figure 2.**
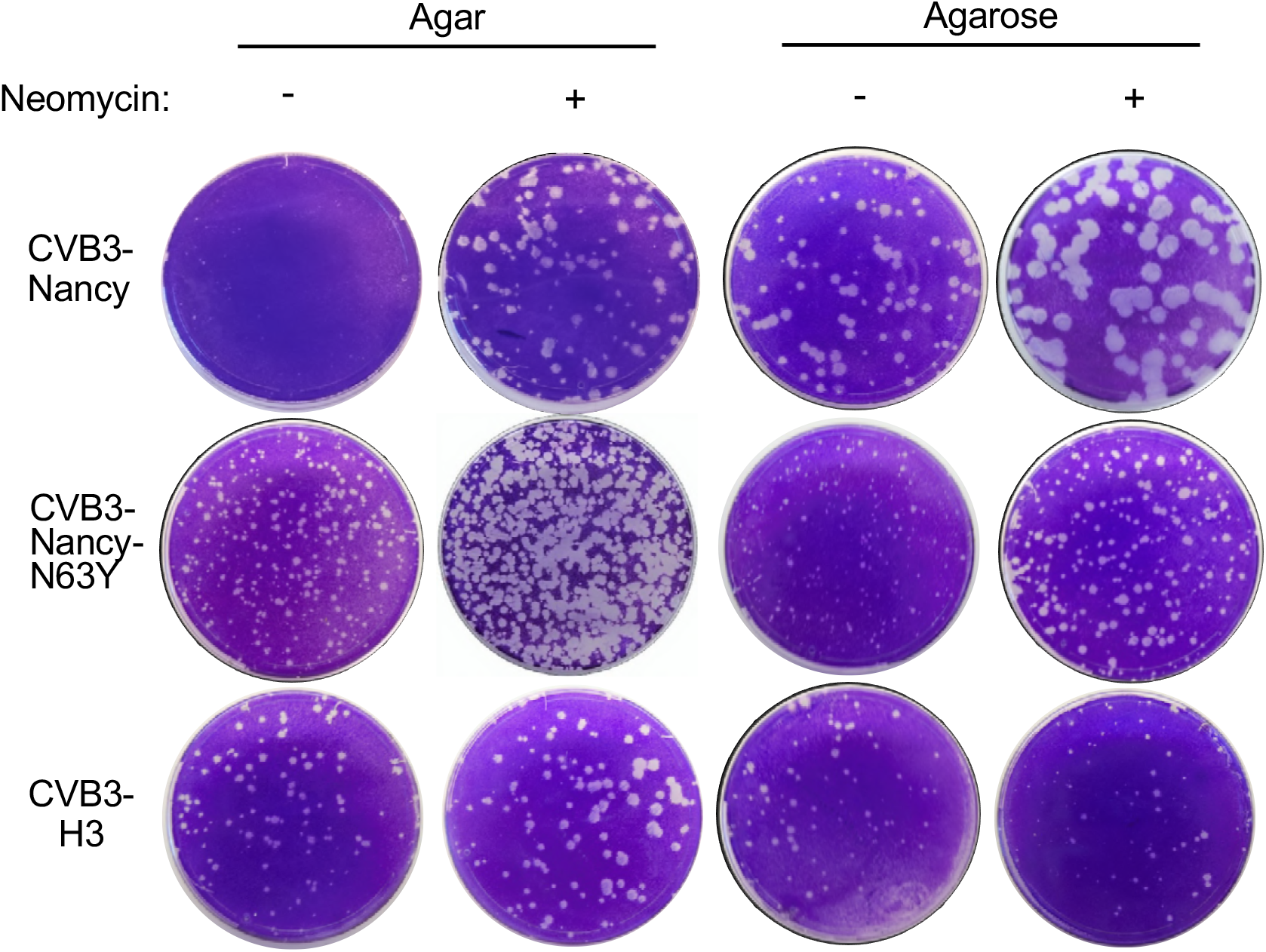
Effect of neomycin on plaque formation of different CVB3 strains. HeLa cells were pretreated with or without neomycin and were infected with approximately 100 PFU of CVB3-Nancy, CVB3-Nancy-N63Y, or CVB3-H3, and then agar or agarose overlays, containing or lacking 1 mg/ml neomycin, were added.

The small plaque size of CVB3-Nancy under agar overlays has been attributed to its binding to sulfated glycans present in agar, which limits viral diffusion (17). However, in agarose overlays, which contain low levels of sulfated glycans, CVB3-Nancy plaque size is significantly larger. Therefore, we sought to examine whether neomycin treatment can increase the plaque size of CVB3-Nancy, CVB3-Nancy-N63Y, and CVB3-H3 in agarose overlays. We found that neomycin increased the plaque size of CVB3-Nancy in presence of an agarose overlay (Fig 2), although the effect was diminished due to larger CVB3-Nancy plaques in agarose overlays. Similarly, CVB3-Nancy-N63Y plaques were slightly larger in the presence of neomycin (Fig 2). However, in agarose overlays, plaque size CVB3-H3 was unaffected by neomycin treatment (Fig. 2). Overall, these data suggest that in both agar and agarose overlays, neomycin can increase the plaque size of CVB3-Nancy and CVB3-Nancy-N63Y, but not of CVB3-H3.

### Early stages of CVB3-Nancy infection are unaffected by neomycin

Given that CVB3-Nancy generates large plaques in the presence of neomycin, we hypothesized that neomycin enhances viral replication. To test this hypothesis, we first determined whether early steps of the viral replication cycle are affected by neomycin. To examine viral attachment, we quantified binding of radiolabeled ^35^S-labeled CVB3-Nancy to HeLa cells in the presence or absence of neomycin pre-treatment. HeLa cells were pre-treated with or without neomycin overnight, followed by incubation with ^35^S-labeled virus at 4°C for 20 min. After washing, cell-associated ^35^S was quantified. ^35^S counts were the same for cells treated with or without neomycin, suggesting that neomycin does not affect viral attachment (Fig. 3A). We next sought to determine if viral translation is affected by neomycin treatment. Picornaviruses, including CVB3, initiate translation early in the viral life cycle, which results in shutoff of host protein synthesis (19, 20).

**Figure 3.**
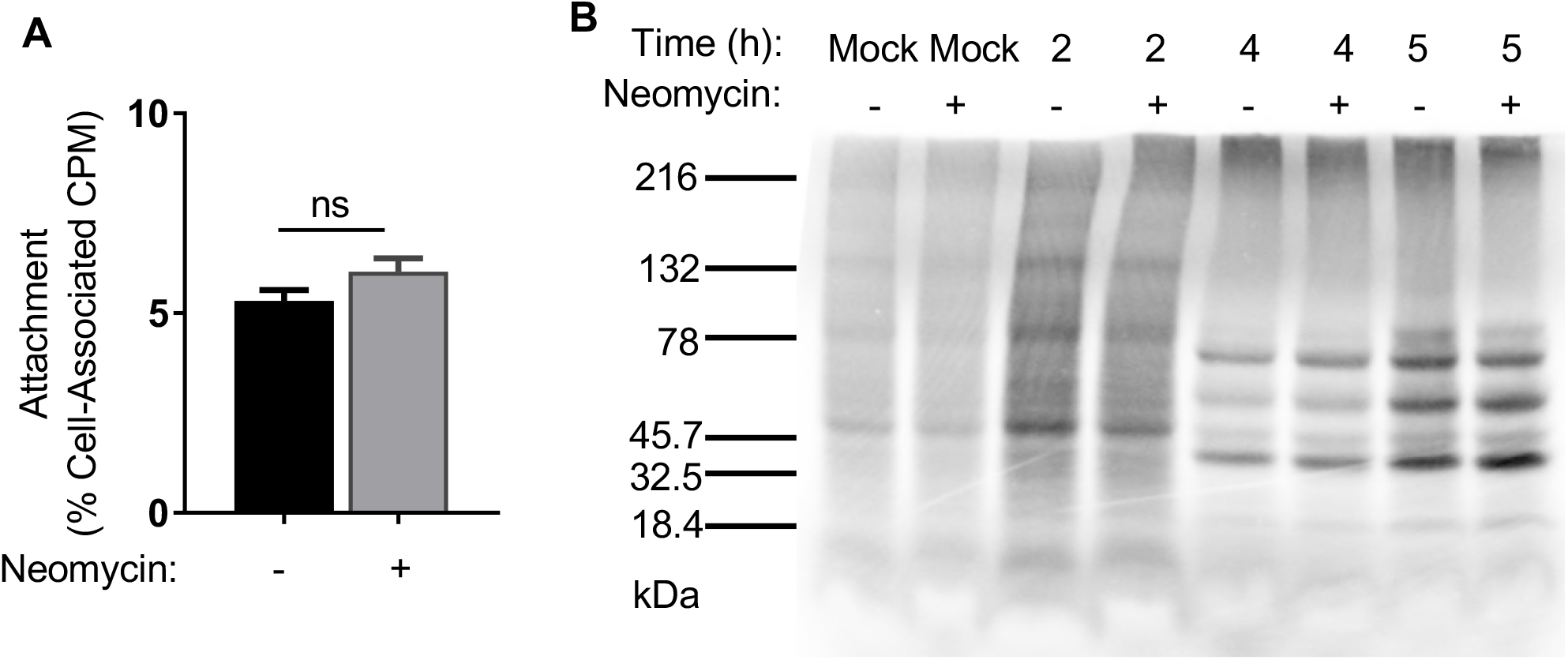
Effects of neomycin on early stages of the CVB3-Nancy replication cycle. A) Cell attachment assay. 1 × 10^6^ HeLa cells were pretreated with or without neomycin prior to incubation with 6000 CPM (6 × 10^6^ PFU) of ^35^S-labeled CVB3-Nancy or no virus (Mock) at 4°C for 20 min to promote viral binding. Cells were washed and ^35^S was quantified in a scintillation counter. Data are means ± standard errors of the means, ns= not significant (unpaired two-tailed Student *t* tests). B) Viral protein synthesis assay. 2.5 × 10^6^ HeLa cells that were pretreated with or without neomycin were inoculated with CVB3-Nancy at an MOI of 20 for 30 min at 37°C. At indicated time points, cells were washed and incubated in 1ml of DMEM lacking methionine and cysteine supplemented with ^35^S-L-methionine and ^35^S-L-cysteine for 15 min at 37°. Cell lysates from equal cell numbers were analyzed on SDS-PAGE gel. Radiolabeled proteins were visualized using a Phosphorlmager.

HeLa cells that were pretreated overnight with or without neomycin were infected with CVB3-Nancy and then at various timepoints cells were exposed to media containing ^35^S-labeled cysteine and methionine to label nascent proteins. Cell lysates were run on an SDS-PAGE gel and ^35^S labeled proteins were imaged via phosphorimager. We found that the amount of labeled viral and cellular proteins were the same for cells treated with or without neomycin (Fig. 3B). Overall, these data suggest that neomycin does not affect the early stages of the CVB3-Nancy life cycle.

### Replication kinetics of CVB3-Nancy are unaffected by neomycin treatment

To examine whether neomycin affects the replication kinetics of CVB3-Nancy, we used single-cycle growth curve assays. HeLa cells that were pretreated with or without neomycin were infected with CVB3-Nancy at an MOI of 0.01, and cell-associated viral titers were determined over time. CVB3-Nancy titers were the same for cells treated with or without neomycin at each timepoint analyzed (Fig. 4A). We next determined whether the presence of an agar overlay could alter the replication kinetics of CVB3-Nancy, and whether the presence of neomycin in the agar overlay could alter CVB3-Nancy growth. For these experiments, HeLa cells were pretreated with or without neomycin, cells were infected for 30 min with CVB3-Nancy at an MOI of 0.01, followed by addition of an agar overlay with or without neomycin. At 0, 2, 4, 6, or 8 hours post-infection, agar overlays were removed, cells were harvested, and cell-associated virus titers were determined by plaque assay on naïve cells. We found no difference in CVB3-Nancy replication in the presence of an agar overlay, with or without neomycin (Fig. 4B). Given that neomycin did not affect single cycle replication kinetics of CVB3-Nancy in either liquid media or agar overlay, we next determined whether neomycin could alter spread during multiple replication cycles. One million HeLa cells were inoculated with 100 PFU of CVB3-Nancy, in the presence or absence of neomycin, and the cells were incubated for 24 h to allow multiple replication cycles to occur. Viral titers in the presence of liquid media showed no differences in viral yield with or without neomycin (Fig. 4C). However, when CVB3-Nancy was grown for 24 h in the presence of an agar overlay, a significant increase in viral yield was detected in the presence of neomycin (Fig. 4D). Overall these data indicate that a single cycle of replication of CVB3-Nancy was unaffected by neomycin treatment, but titers from multiple cycles were increased by neomycin treatment when agar overlays were present, suggesting neomycin may aid viral spread in the presence of agar.

**Figure 4.**
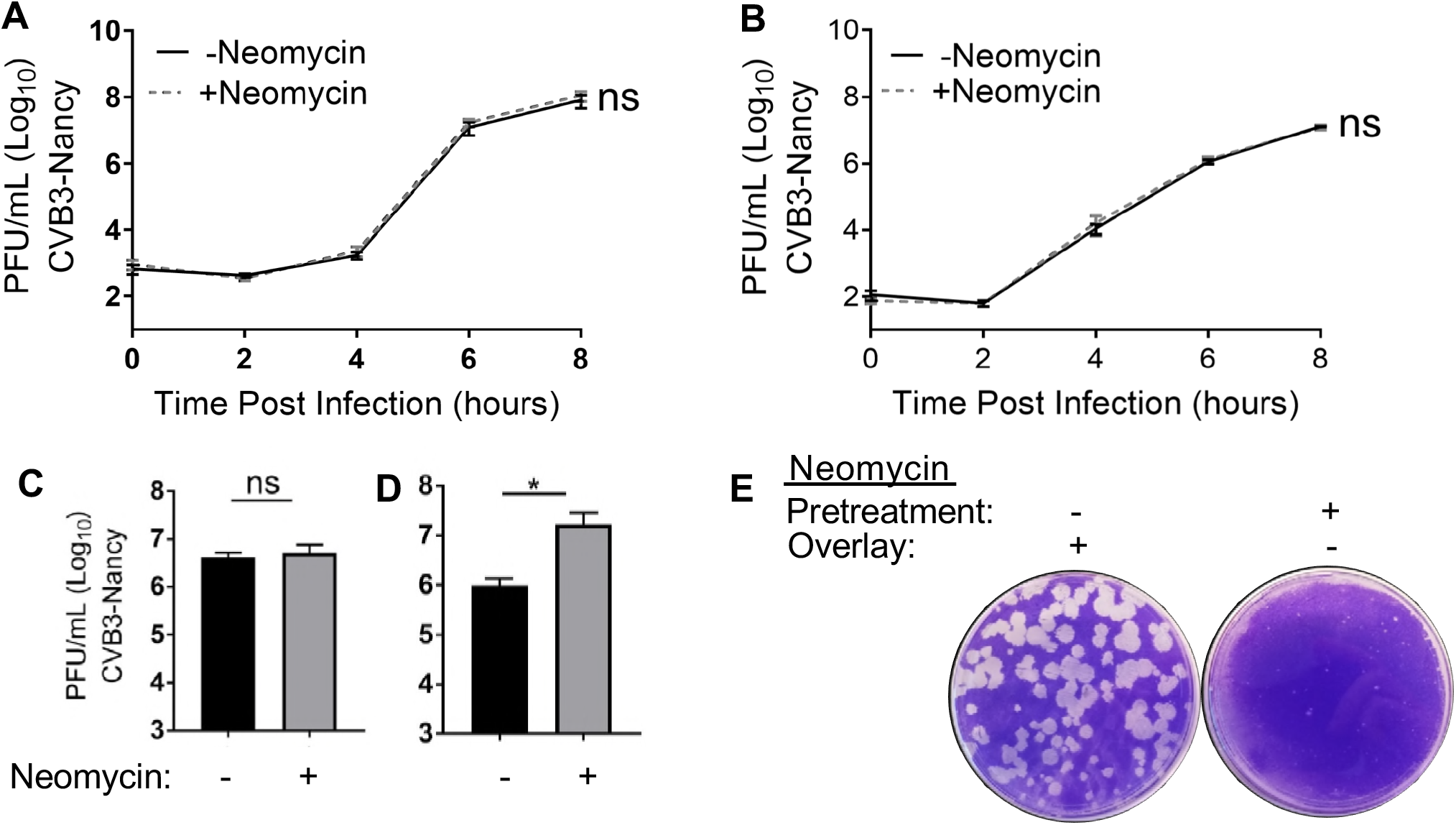
Effects of neomycin on CVB3-Nancy replication kinetics. Growth curve in presence of liquid media (A) or agar overlay (B). Briefly, 1 × 10^6^ HeLa cells that were pretreated with or without neomycin were inoculated with CVB3-Nancy at an MOI of 0.01. Virus was incubated for 30 min at 37°C and either DMEM liquid (A) or 1% Agar/1% DMEM mixture (B) with or without neomycin was added. At indicated timepoints, intracellular virus was harvested and quantified by plaque assay. To examine multi-cycle replication and spread, cells were infected with 100 PFU of CVB3-Nancy and either DMEM (C) or 1% Agar/1% DMEM mixture (D) with or without neomycin followed by plaque assay of cell-associated virus. E) HeLa cells were pretreated with or without 1 mg/ml of neomycin prior to plating 100 PFU of CVB3-Nancy on cells with agar overlays with or without 1 mg/ml of neomycin. Plates were stained with crystal violet 48 hpi. *, P<0.05 (unpaired two-tailed Student *t* tests), ns= not significant.

### Positive charge of neomycin contributes to generation of large CVB3-Nancy plaques

Because neomycin increased plaque size of CVB3-Nancy in agar overlays and neomycin increased 24 h titers of CVB3-Nancy when an agar overlay was present, we hypothesized that neomycin enhances viral diffusion by overcoming inhibition by negatively charged compounds in agar overlays. Additionally, we hypothesized that neomycin must be present in the agar overlay to enhance plaque formation and that pre-treatment of cells with neomycin would not be sufficient to increase plaque size when neomycin was not present in agar overlays. To test this, we pretreated cells with or without neomycin, infected with 100 PFU of CVB3-Nancy and added an agar overlay with or without neomycin. We found that large plaques formed in cells with neomycin in agar overlays regardless of whether cells were pretreated with neomycin before infection. Conversely, small plaques formed in cells without neomycin in agar overlays regardless of whether they were pretreated with neomycin (Fig. 4E). Thus, the presence of neomycin in the overlay is sufficient to increase plaque size of CVB3-Nancy.

Agar is rich in anionic sulfated polysaccharides, which inhibits some viruses by binding and preventing cell adsorption or diffusion (21–23), and cationic compounds can overcome this negative charge inhibition of the agar overlay (22). Given that neomycin is also a positively charged compound (24), we hypothesized that neomycin increases CVB3-Nancy plaque size by overcoming the inhibitory negative charge of the agar overlay. To test this, we evaluated whether other positively charged compounds could also increase the plaque size of CVB3-Nancy. We found that two positively charged compounds, poly-L-lysine and protamine, increased plaque size of CVB3-Nancy (Fig. 5A). We next determined whether neomycin was capable of overcoming the inhibitory effect of negatively-charged heparin (17). We found that CVB3-Nancy was able to generate large plaques in the presence of heparin when neomycin was present (Fig. 5B). These data indicate that the positive charge of neomycin contributes to large plaque formation of CVB3-Nancy.

**Figure 5.**
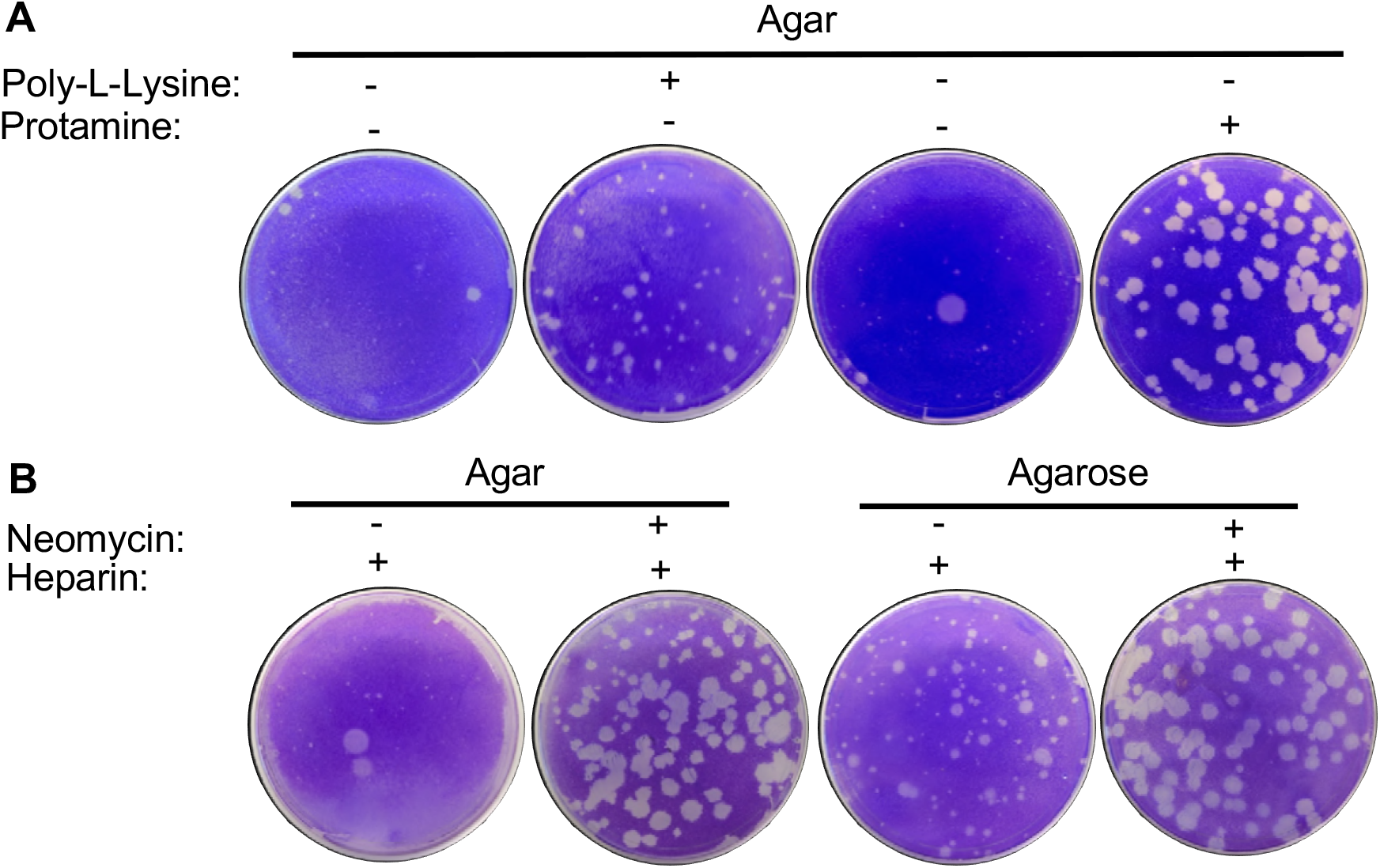
Effect of positively or negatively charged compounds on CVB3-Nancy plaque formation. A) 8.8 × 10^6^ HeLa cells were infected with 100 PFU of CVB3-Nancy and agar overlays were added with or without 0.1μM poly-L-lysine or 0.8 mg/ml protamine (positively charged compounds). B) 8.8 × 10^6^ HeLa cells were infected with 100 PFU of CVB3-Nancy and neomycin-containing agar or agarose overlays were added with or without 1 mg/ml of heparin (negatively charged compound).

Because other positively charged compounds phenocopy neomycin’s effects, we hypothesized that neomycin facilitates large plaque formation of CVB3-Nancy by neutralizing negatively charged inhibitory molecules in agar, allowing the virus to diffuse more efficiently. To test this, we used a diffusion assay previously described by Wallis and Melnick (22). Agar overlays with or without neomycin or protamine were added to untreated and uninfected HeLa cells and overlays were allowed to solidify. Then, 5 × 10^4^ PFU of CVB3-Nancy was added dropwise on top of the agar overlay. Virus was allowed to diffuse downward through the ~1 cm-thick agar overlay and infect cells and plates were stained with crystal violet at 1, 2, or 3 dpi to examine cell death. We found that CVB3-Nancy diffusion and subsequent cell death was greatly enhanced when either neomycin or protamine was added to the agar overlays, when compared to agar overlays with no treatment (Fig. 6). Overall these results suggest that the positive charge of neomycin enhances CVB3-Nancy diffusion by overcoming the negative charge of agar overlays.

**Figure 6.**
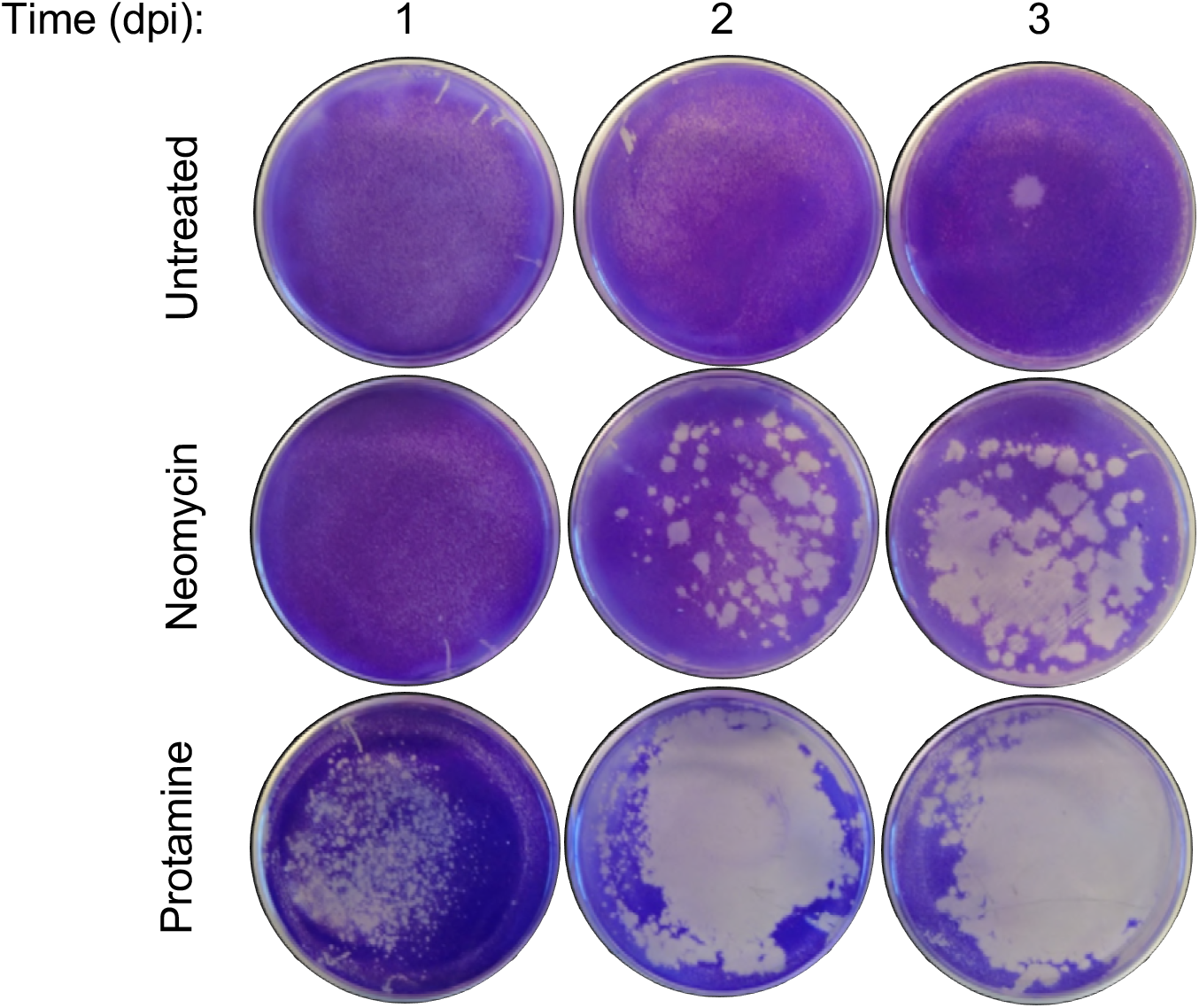
Effects of positively charged compounds on CVB3-Nancy diffusion. 1 × 10^6^ HeLa cells in 60 mm tissue culture plates were overlaid with 4 ml of 1% Agar/1% DMEM mixture that contained or lacked 1 mg/ml neomycin or 0.8 mg/ml protamine. Once the overlay had solidified, 5 × 10^4^ PFU of CVB3-Nancy in 200 μL was added dropwise to the top of the overlay. Cells were placed at 37°C to allow diffusion of the virus through the overlay to the cell monolayer and plates were stained with crystal violet at 1, 2, or 3 days post infection (dpi) to reveal the extent of cell death from viral replication.

## Discussion

Although it is known that antibiotics deplete the gut microbiota of mice and microbiota depletion reduces infection of certain enteric viruses, it remains unclear if antibiotics can directly affect enteric viruses independently of effects on the microbiota. Here we shown that an aminoglycoside antibiotic, neomycin, increases plaque size of CVB3-Nancy. In this study, we found that neomycin enhances CVB3-Nancy plaque formation through increased viral diffusion due to its positive charge.

CVB3-Nancy is a cell culture adapted virus that has increased binding to heparan sulfate, a negatively charged sulfated polysaccharide that is present on the surface of cells and in agar overlays (17, 21). The presence of sulfated polysaccharides in agar overlays limits diffusion of certain heparan sulfate binding viruses, which results in formation of small plaques (21). Interestingly, CVB3-Nancy-N63Y, a derivative of CVB3-Nancy that contains a single mutation that decreases virion binding to sulfated polysaccharides, also had increased plaque size due to neomycin treatment. CVB3-Nancy-N63Y has larger plaques in agar overlays, suggesting its diffusion is less limited by the negative charge of the sulfated polysaccharides (Fig. 2). CVB3-H3 is a less culture adapted and more pathogenic strain of CVB3, and CVB3-H3 formed large plaques either in presence or absence of neomycin (Fig. 2). Since neomycin only had an effect on plaque formation of CVB3-Nancy strains, but not CVB3-H3 or poliovirus, neomycin may only enhance diffusion of viruses that bind to negatively charged sulfated polysaccharides.

Anionic polymers, such as sulfated polysaccharides present in agar overlays, have inhibitory effects by limiting adsorption of newly formed virions to cells (22, 25, 26). Protamine, a positively charged compound, enhances plaque size of encephalomyocarditis virus and adenovirus (22, 25, 26), promotes diffusion of enterovirus (27), and enhances infectivity of rabies virus (28). In agreement with previous studies, we found that protamine significantly enhanced plaque size of CVB3-Nancy in the presence of an agar overlay (Fig 5A). We also found that poly-L-lysine, a compound with similar charge to neomycin (29), also increased CVB3-Nancy plaque size in the presence of an agar overlay. To confirm that the positive charge of neomycin enhances viral diffusion, we performed a diffusion assay and found that neomycin or protamine treated overlays enhanced CVB3-Nancy diffusion.

In conclusion, we found that positively charged compounds, such as neomycin, poly-L-lysine, and protamine, aid CVB3-Nancy and CVB3-N63Y diffusion in the presence of either an agar or agarose overlay. This work provides insight into methods to enhance plaque formation and reveals that a commonly used antibiotic can have microbiota-independent effects on a virus.

## MATERIALS AND METHODS

### Cells and virus

HeLa cells were grown in Dulbecco’s modified Eagle’s Medium (DMEM) supplemented with 10% calf serum (Sigma-Aldrich) and 1% penicillin-streptomycin (Sigma-Aldrich). The CVB3-Nancy and CVB3-H3 infectious clones were obtained from Marco Vignuzzi (Pasteur Institute, Paris, France) and the CVB3-Nancy-N63Y infectious clone was previously generated from CVB3-Nancy by site directed mutagenesis (17). The poliovirus infectious clone was serotype 1 Mahoney (30). Viral stocks were prepared as previously described (17) and viral titers were determined by plaque assay as previous described (17, 31). Briefly, monolayers of HeLa cells were infected for 30 min followed by addition of an overlay containing media and 1% agar (Becton Dickinson) or 1% SeaKem™ LE Agarose (Lonza). Following incubation, plaques were visualized by staining with an alcholic solution of crystal violet. Neomycin-treated cells were pretreated overnight with 1 mg/ml of neomycin (Research Products International).

### CVB3 cell attachment

^35^S-labeled CVB3-Nancy was generated as previously described (8)(17). Briefly, cells were infected with CVB3-Nancy in the presence of ^35^S-L-methionine and ^35^S-L-cysteine Express Labeling Mix (PerkinElmer), and viruses in cell lysates were purified using CsCl gradient ultracentrifugation (8)(17). 6000 CPM (6 × 10^6^ PFU) was incubated with 1 × 10^6^ HeLa cells that were pretreated with or without 1 mg/ml neomycin at 4°C for 20 min to promote viral binding. Cells were washed three times with ice cold phosphate-buffered saline (PBS) to remove unbound labeled virus, trypsinized, and ^35^S was quantified in a scintillation counter (Beckman Coulter, LS6500 Multi-Purpose Scintillation Counter).

### CVB3 translation assay

2.5 × 10^6^ HeLa cells were pretreated with or without 1 mg/ml of neomycin for 16 h and were inoculated with CVB3-Nancy at an MOI of 20 for 30 min at 37°C. Inoculum was aspirated, cells were washed with PBS, and complete DMEM was added. At 2, 4, and 5 hours post infection (hpi), cells were washed and incubated in 1 ml of DMEM lacking methionine and cysteine (Sigma-Aldrich) with 55 μCi of ^35^S Express Labeling Mix (PerkinElmer) for 15 min at 37°. Cells were harvested and lysed in buffer containing 10 mM Tris pH 8, 10 mM NaCl, 1.5 mM MgCl2, and 1% NP-40, and nuclei were removed by centrifugation. Supernatants from equal cell numbers were analyzed on 4–20% Mini-PROTEAN^®^ TGX™ Precast Protein Gels (Bio-Rad). Gels were dried at 80°C for 1 hour and exposed to a phosphorimager screen overnight. Radiolabeled proteins were visualized using a Phosphorlmager (Typhoon FLA 9500).

### Viral growth curves

1 × 10^6^ HeLa cells pretreated overnight with or without 1 mg/ml neomycin were inoculated with CVB3-Nancy at an MOI of 0.01. Virus was incubated for 30 min at 37°C to promote viral binding, inoculum was aspirated, the cell monolayer was washed once in PBS, and 2 ml of either DMEM or 1% Agar/1% DMEM mixture with or without 1 mg/ml neomycin was added. At 0, 2, 4, 6, and 8 hours post infection, media was removed, the cell monolayer was washed once in PBS, and cells were trypsinized and pelleted. Intracellular virus was harvested by freeze-thawing three times. For 24 hpi assays, cells were infected with 100 PFU of CVB3-Nancy and 2 ml of either DMEM or 1% Agar/1% DMEM mixture with or without 1 mg/ml neomycin was added. Plaque assay, as described above, was used to quantify amount of intracellular PFU.

### Viral diffusion assay

1 × 10^6^ uninoculated HeLa cells in 60 mm tissue culture plates were overlaid with 4 ml of 1% Agar/1% DMEM mixture with or without 1 mg/ml neomycin or 0.8 mg/ml protamine. Once the overlay had solidified, 5 × 10^4^ PFU of CVB3-Nancy in 200 μL was added dropwise to the overlay. Cells were placed at 37°C to allow diffusion of the virus through the overlay to the cell monolayer. Plates were stained with crystal violet at 1, 2, or 3 days post-infection.

### Statistical analysis

The difference between groups were examined by unpaired two-tailed Students t-test. Error bars represent the mean ± the standard error of the mean. P<0.05 was considered significant. All analysis of data were performed using Graph Pad Prism version 7.00 for Windows, GraphPad Software, La Jolla California USA.

## Acknowledgements

We thank Andrea Erickson and Broc McCune for helpful comments on the manuscript.

## Funding Information

Work in J.K.P.’s lab is funded through NIH NIAID grant R01 AI74668, a Burroughs Wellcome Fund Investigators in the Pathogenesis of Infectious Diseases Award, and a Faculty Scholar grant from the Howard Hughes Medical Institute. MWA was supported in part by NIH NIAID grant T32 AI007520.

